# The Effect of a Fennel Extract on the STAT Signaling and Intestinal Barrier Function

**DOI:** 10.1101/2021.07.01.450766

**Authors:** John Rabalais, Philip Kozan, Tina Lu, Nassim Durali, Kevin Okamoto, Barun Das, Matthew D. McGeough, Jae Beom Lee, Kim E. Barrett, Ronald Marchelletta, Mamata Sivagnanam

**Author notes:** Corresponding Author: Mamata Sivagnanam, Division of Gastroenterology, Hepatology and Nutrition, Department of Pediatrics, 9500 Gilman Drive, La Jolla, CA 92093 USA, University of California San Diego, Rady Children’s Hospital San Diego.

## Abstract

**Background:** *Foeniculum vulgare*, *F. vulgare*, commonly known as fennel, is believed to be one of the world’s oldest medicinal herbs and has been exploited by people for centuries as a nutritional aid for digestive disorders. In many southeast Asian countries it is ingested as an after-meal snack, mukhvas, due to its breath-freshening and digestive aid properties. *F. vulgare* is used in some countries, such as Iran, as a complementary and alternative treatment for inflammatory bowel disease (IBD).

**Methods:** This study investigated the effects of *F. vulgare* on the barrier function of the intestinal epithelium Signal Transducer and Activator of Transcription (STAT) pathway, which is active in inflammatory bowel disease.

To study the protective effects of *F. vulgare* extract *in vitro*, monolayers derived from the T84 colonic cell line were challenged with interferon-gamma (IFN-γ) and monitored with and without *F. vulgare* extract. To complement our *in vitro* studies, the dextran sodium sulfate induced murine colitis model was employed to ascertain whether the protective effect of *F. vulgare* extract can be recapitulated *in vivo*.

**Results:** *F. vulgare* extract was shown to exert a protective effect on TEER in both T84 and murine models and showed increases in tight junction-associated mRNA in T84 cell monolayers. Both models demonstrated significant decreases in phosphorylated STAT1 (pSTAT1), indicating reduced activation of the STAT pathway. Additionally, mice treated with *F. vulgare* showed significantly lower ulcer indices than control mice.

**Conclusions:** We conclude barrier function of the gastrointestinal tract is improved by *F. vulgare*, suggesting the potential utility of this agent as an alternative or adjunctive therapy in IBD.

## Introduction

*Foeniculum vulgare* (*F. vulgare*), commonly known as fennel, is a flowering plant species in the family Apiaceae. It is believed to be one of the world’s oldest medicinal herbs and has been exploited by people for centuries for its reported anti-inflammatory and antipathogenic properties.[1] In many southeast Asian countries it is ingested as an after-meal snack, mukhvas, due to its breath-freshening and digestive aid properties.

There is a rising interest in herbal therapies for IBD worldwide, with clinical studies being done on a variety of natural products, such as *aloe vera* gel and Andrographis paniculate extract, which proved effective compared to placebos.[2] Previous studies have examined various products derived from *F.vulgare*, though not directly in the intestine. One study showed that oil extracted from *F.vulgare* was characterized to have safe antithrombotic activity through its broad spectrum antiplatelet activity, clot destabilizing effect, and vasorelaxant action.[3] This study also demonstrated that fennel oil provided significant protection from ethanol-induced gastric lesions in rats. Fennel oil extract has been shown to have *in vitro* antifungal activity when column fractions are screened against MDR strains of *Mycobacterium tuberculosis*.[4] Anethole, a major component of fennel oil, is an organic compound that is widely used as a flavoring substance and also has been shown to have anti-proliferative effects on prostate cancer cells.[5] It is a derivative of phenylpropene, a type of aromatic compound that occurs widely in nature. In addition to the beneficial effects of fennel oil, fennel “waste” (components remaining after oil extraction) has been studied and found to exhibit high antioxidant activity. [6] [7] *F. vulgare* is also used in some countries, such as Iran, as a complementary and alternative treatment for inflammatory bowel disease.[1, 8, 9]. Additionally, a combination of *F. vulgare* and turmeric oils has been shown to increase quality of life in IBS patients, highlighting the wide-ranging benefits of *F. vulgare* on the body, particularly in the gastrointestinal system. [10] Other herbal therapy studies have shown far reaching effects in the context of IBD, with data suggesting herbal remedies are able to mediate damage to barrier function, immune response, and even restore gut microbiota.[11] This study specifically investigates the use of *F. vulgare* in mediating mechanisms of inflammatory bowel disease.

Inflammatory bowel disease affects nearly 1 million people annually in the United States.[12] IBD is characterized by chronic inflammation of the small and/or large intestines. The pathogenesis of IBD involves a multitude of genetic and environmental factors that are presumed to cause an excessive and inappropriate mucosal immune response.[12–14] In IBD, the balance between pro- and anti-inflammatory mediators is shifted, leading to infiltration of the lamina propria with immune cells that release pro-inflammatory cytokines such as interferon-gamma (IFN-γ). To ensure intestinal homeostasis, a robust barrier between epithelial cells is essential to protect against foreign antigens. The barrier formed by epithelial cells is partly comprised of the apically-located tight junction complex, which includes occludin (OCLD), claudins, and tight junction protein-1 (TJTP-1).[15] In IBD, this barrier becomes compromised.[16] The cytokines present in IBD damage the intestinal barrier, resulting in clinical and pathologic manifestations including mucosal friability, decreased tissue resistance, and increased paracellular permeability.[17]

IFN-γ is known to signal through the Janus Kinase (JAK)/ Signal Transducer and Activator of Transcription (STAT) pathway, and activation of this pathway, including phosphorylation of STAT1, has been demonstrated in IBD.[18, 19] The pathway is initiated when IFN-γ binds to its receptor and induces dimerization of its subunits. The JAK dimer is then recruited to the receptor and becomes activated, phosphorylating STAT proteins.[20–22] *In vitro* and *in vivo* studies show that activated IFN-γ receptor can modify epithelial barrier function by mechanisms that include new protein synthesis, membrane trafficking, kinase activation, cytoskeletal modulation and epithelial apoptosis.[23] Aberrant activation of IFN γ receptor induces epithelial dysfunction in a similar pathology observed in Crohn’s Disease.[18, 24, 25] Of interest, the JAK/STAT pathway is targeted by Tofacitinib, a novel small-molecule drug investigated for its ability to inhibit the JAK family in the setting of IBD. This results in immunosuppression through downregulation of inflammatory molecules [26, 27] Previous data have shown that components of *F. vulgare* oil extract potently inhibit TNF-induced activation of NF-kB in B cells[28] and also inhibit the release of interleukin-1β following administration of LPS to rats.[29] Based on the known impact of *F. vulgare* on responses evoked by other inflammatory mediators, we hypothesized that *F. vulgare* may also play a role in modulating the JAK/STAT pathway. The role of *F. vulgare* on intestinal barrier function and the JAK/STAT pathway has yet to be studied and could provide an additional mechanism underlying *F. vulgare’s* anti-inflammatory properties and its use as a complementary treatment for IBD. In this study, we utilize the T84 cell line to examine any protective effects of *F. vulgare* in cells treated with IFN-γ. This colonic adenocarcinoma cell line has been widely used as a model for studies of epithelial barrier function.[30] To understand how *F. vulgare* may ameliorate IBD, a murine model of dextran sodium sulfate (DSS)-induced colitis was employed.[31]

## Methods

### *F. vulgare* extract

*F. vulgare* extract was obtained from Nature’s Answer (Hauppauge, NY) and was produced by crude extraction from dry *F. vulgare* seeds by the company. Ingredients found in the extract include: Fennel seed extract, 15% Certified Organic Ethyl Alcohol, water, and Vegetable Glycerin. Nature’s Answer performs DNA based quality control to ensure the presence of fennel seed within the extract in addition to other appropriate quality control processes recommended by the FDA.

### Cell culture

T84 cells were cultured in 1:1 Dulbecco’s modified Eagle’s Medium/F-12 Ham’s medium with 15 mM L-glutamine (Mediatech Inc., Manassas, VA), 5% bovine calf serum (Invitrogen, Grand Island, NY), and 1% penicillin-streptomycin (Mediatech Inc.) and maintained according to a standard protocol.[32] Cells were divided into 6 groups: control (receiving ethanol as vehicle), IFN (vehicle and 100 U/mL IFN-γ), and *F. vulgare/IFN* treatments with 4 increasing dosages of *F. vulgare* extract (4.5, 6, 7.5 and 9 μL/mL) with IFN (100 U/mL IFN-γ). *F. vulgare* treatments began 2 days after culturing on 24 well plates and IFN was added 2 days later. Groups were plated in triplicate at 0.5 x 10^6^ cells per well.

RNA and protein were collected 72 hours after IFN treatment. To eliminate the concern of *F. vulgare* extract toxicity on T84 cells, increasing concentrations of *F. vulgare* extract were diluted in cell media. Epithelial health was monitored using barrier function as a readout measured by transepithelial electrical resistance (TEER).[33]

### Cell Viability Analysis

Cell viability of the T84 cells were assessed with propidium iodide (PI) staining using flow cytometry in a similar way as described earlier [34] after treatment with Fennel extract and vehicle (Ethanol). The single cell suspension of T84 cells were obtained after trypsinization for 5 minutes at 37°C followed by mechanical disruption and passing through 70 micron cell strainer (Corning #352235, Corning, NY) to separate cell clumps. The cells were resuspended in PBS and PI (Cat# P4864, Sigma-Aldrich, St Louis, MO) was added to the cell suspension at 0.5 μg/mL. After incubation for 15 minutes in the dark, the cells were acquired in BD Accuri^™^ C6 (BD, Franklin lakes, NJ) Flowcytometer. The percentage of PI positive cells were analyzed with FlowJo software (FlowJo LLC, Ashland, OR).

Among the different concentrations of *F. vulgare* extract used, the highest concentration (9 μL/mL) was chosen to check for toxicity in T84 cells. The vehicle treated group was also included in the experiment to assess any negative effects of ethanol. Forward scatter (FSC)-Area (A) vs FSC-Height (H) pulse gating strategy was incorporated to exclude the duplets and cell aggregates from the total cell population as described earlier [34]. The PI positive staining in the gated population was calculated after normalizing the mean fluorescent intensity of the respective unstained population.

### Murine colitis studies

C57BL/6 mice obtained from Jackson Labs (Bar Harbor, ME) were used for all *in vivo* studies. Mice weighed 20-25 grams and were housed in plastic cages with flake bedding. Mice were randomized to 4 groups. All mouse groups were orally gavaged (150 μL) daily for 8 days with water (control and DSS group), or *F. vulgare* extract dissolved in drinking water at 4.5 uL/mL (*F. vulgare* alone, DSS/*F. vulgare*), a concentration used in previous murine studies during the daytime.[3, 35] DSS and DSS/*F. vulgare* groups received 3.5% DSS in their drinking water for 5 days followed by 3 days of normal drinking water in their group cages. Mice had access to chow and water and rooms were kept on light dark cycle. Mice were euthanized per UCSD protocol using carbon dioxide. All studies were approved by the UCSD Institutional Animal Care and Use Committee and no adverse events were noted. Upon sacrifice of mice, gross health of colonic tissue (friability) and stool characteristics, such as presence of blood, color, softness, etc., were noted.

### Electrical resistance

For resistance studies, T84 cells were grown on semipermeable 12 mm Millicell-HA cell culture inserts with 0.5×10^6^ cells added per insert. Cells were maintained in media as above. Cell culture media was changed every 2 days for approximately 10 days until a stable monolayer was established.[32] To assess monolayer integrity TEER was measured across T84 monolayers with a voltohmeter (World Precision Instruments, Sarasota, FL). The monolayer was considered mature when the TEER of the monolayer reached a stable value of approximately 1000 ohms.cm^2^. In *F. vulgare* pre-treatment studies, *F. vulgare* extract or vehicle was added to the cells upon reaching this baseline. TEER values continued to be taken for each of the groups. After 2 days, IFN-γ was added basolaterally to the IFN and *F. vulgare* groups and TEER values were followed for 5 days. In *F. vulgare* rescue studies, cells were plated in the same manner as pre-treatment studies. Upon reaching mature monolayer, either IFN-γ or vehicle was given to corresponding groups and *F. vulgare* was added to monolayers 2 days post IFN-γ or vehicle addition.

For murine studies, animals were sacrificed by carbon dioxide exposure per institutional guidelines and full-thickness segments of mid colon were mounted in Ussing chambers (Physiological Instruments, San Diego, CA) per protocol (mouse tissue window area: 0.09 cm^2^).[36] Colonic tissue was excised and washed in Ringer’s solution containing 140 mM Na^+^, 5.2 mM K^+^, 1.2 mM Ca^2+^, 0.8 mM Mg^+^, 120 mM Cl^-^, 25 mM HCO_3_^-^, 2.4 mM H_2_PO_4-_, 0.4 mM HPO_4_^2-^, 10 mM glucose. Tissue was then mounted into the slides and bathed in an oxygenated Ringer’s solution at 37°C that had been previously equilibrated in the Ussing chambers for 30 minutes. A transepithelial current pulse of 1 μA was administered through the chamber and the resulting voltage drop between chambers was measured to assess tissue viability and TEER was calculated from the resulting voltage deflection using Ohm’s law (V=IR).

### Western blotting

T84 cells were suspended in RIPA lysis buffer and proteins were extracted and analyzed according to a published protocol.[37] Samples were resuspended in loading buffer (50 mM Tris (pH 6.8), 2% SDS, 100 mM dithiothreitol, 0.2% bromphenol blue, and 20% glycerol).

Mid colon samples were placed in nonyl phenoxypolyethoxyethanol-40 (NP-40) buffer containing 0.9% NaCl, 10% glycerol, 50 mM Tris (pH 2.8), 0.1% NP40, 5 mM EDTA, 20 μM NaF, 1 μg/ml antipain, 1 μg/ml pepstatin, 1 μg/ml leupeptin, 1 mM NaVO3, and 100 μg/ml phenylmethylsulfonyl fluoride and lysed using a mini bead beater (BioSpec Products, Bartlesville, OK). The lysate was centrifuged and the supernatant containing the proteins was removed.[37] Samples were suspended in loading buffer as above.

Cell and tissue lysates were diluted at a 1 to 1 ratio of lysate to loading buffer and loaded onto Mini-Protean TGX precast gels (BioRad), electrophoresed, then transferred onto PVDF membranes. Membranes were blocked with 5% bovine serum albumin/TBST.

Western blotting was performed using rabbit antibodies to pSTAT1 and STAT1 (Cell Signaling Technologies, Beverly, MA) at 1:1000. A mouse monoclonal antibody to β-actin at 1:5000 (Sigma) was used to correct for loading. Horseradish peroxidase-conjugated anti-mouse or anti-rabbit IgG secondary antibodies (Cell Signaling Technologies, Beverly, MA) were used at 1:2000 dilutions. A semiquantitative measurement of band density was performed using ImageJ for Windows software.

### Q-PCR

Total RNA from T84 cells was isolated using Direct-zol RNA MiniPrep kits (Zymo, Irvine, California). RNA was extracted initially using TRIzol (Invitrogen, Carlsbad, California). First strand cDNA was synthesized with iScript cDNA Synthesis kit (Bio Rad, Irvine, California) using the recommended protocol. Real time PCR reactions were set up using FastStart Universal SYBR Green Master Mix (Invitrogen) and thermal cycling performed on a StepOnePlus Real-Time PCR System using Step One software v2.0. (Applied Biosystems, Carlsbad, CA). Primers were obtained from IDT (Integrated DNA Technologies, Coralville, IA). Glyceraldehyde 3-phosphate dehydrogenase (GAPDH) was used as the housekeeping gene. STAT-1 and GAPDH primers were diluted to 100μM. TJTP-1 and OCLD primers were diluted to 50μM.

### Tissue Imaging

Segments of mid colon were excised and fixed with 4% paraformaldehyde for 24 hours at room temperature, then paraffin-embedded and sectioned onto glass slides. De-paraffination was performed and samples were heated in 10mM sodium citrate buffer. Hemotoxylin and eosin staining was performed by the Allergy Institute of La Jolla.[38] Imaging was performed using a Leica DMi1 inverted microscope at 20x magnification using LAS 4.10 EZ acquisition software.

### Ulcer indices

To assess the effect of *F. vulgare* on intestinal structural integrity, ulcer indices were assigned on a scale of 0 to 19 based on an established scoring system assessing inflammation on tissue sections[39] by four reviewers blinded to the experimental condition. Criteria for assessment of colonic inflammation and ulceration were as follows:

- Inflammation: 0 – Normal, 1-Minimal infiltration of lamina propria, focal to multifocal, 2 – Mild infiltration of lamina propria, multifocal, mild gland separation, 3 – Moderate to mixed infiltration, multifocal with minimal edema, 4 – Marked mixed infiltration into submucosa and lamina propria with extensive areas of gland separation, enlarged Peyer’s patches, edema
- Epithelium: 0 – Normal, 1 – Minimal damage, focal mucosal hyperplasia, 2 – Mild damage, multifocal tufting of rafts of epithelial cells with increased numbers of goblet cells, 3 – Moderate damage, extensive local or multifocal erosion or epithelial attenuation, 4 – Marked locally extensive mucosal ulceration
- Glands: 0 – Normal, 1 – Rare gland dilation present, 2 – multifocal gland dilation, 3 - multifocal gland dilation with abscessation and occasional loss of glands, 4 – locally extensive to subtotal loss of glands
- Depth of Lesion: 0 – None, 1 – Mucosa, 2 – Mucosa and submucosa, 3 – Transmural
- Extent of Section Affected: 0 – None, 1 – Minimal < 10%, 2 – Mild 10-25%, 3 – Moderate 26-50%, 4 – Severe > 50%

Each of the criteria were given separate scores and summed to give an overall index score per image. Four mice were present in each treatment condition (Vehicle, FN, DSS, FN/DSS). Four random images were taken from each mouse and provided to the reviewers for blinded scoring. The average ulcer index score for each mouse was generated.

### Statistical analysis

One-way and Two-way Anova were performed using GraphPad Prism version 8.00 for Windows (GraphPad Software, La Jolla, CA). Data from all animal experiments were included in data analysis. All raw data used and analyzed during the current study are available from the corresponding author on request.

## Results

### F. vulgare exerts protective effects on barrier integrity in T84 cells

TEER correlates positively with epithelial barrier function.[33] As expected, addition of IFN-γ reduced the TEER of T84 cell monolayers within 1-2 days (p<0.01).[40] However, in T84 cells that were pretreated with *F. vulgare*, there was no significant decrease in TEER upon addition of IFN. The FN-treated cells displayed a 30-50% increase TEER over the course of the treatment, matching the growth rate of vehicle-treated control cells, indicating little to no adverse effects due to IFN-γ treatment (Figure 1A). In order to assess whether the *F. vulgare* treatment has any detrimental effect on T84 cells, a cell viability assay was performed with PI staining using flow cytometry at day 1 and day 5 (maximum length of the experiments) of post treatment. The percentage of PI positive dead cells was found to be equivalent among non-treated group, vehicle and fennel treated group at both day 1 and day 5 post treatment. This result confirmed that both the fennel extract and ethanol vehicle in the said concentration have no detrimental effect on T84 cells viability (Figure 1C). In a separate experiment, *F. vulgare* was administered to T84 monolayers after IFN-γ treatment. As before, IFN-γ addition immediately decreased TEER values, however upon addition of *F. vulgare*, TEER values were able to be stabilized, preventing further loss of barrier integrity, and in two cases restored back to baseline levels (Figure 1B). By day 5 of the experiment, FN 1, FN 3 and FN 4 treatments showed a significant increase in TEER values compared to IFN-*γ* treated cells.

**Figure 1.**
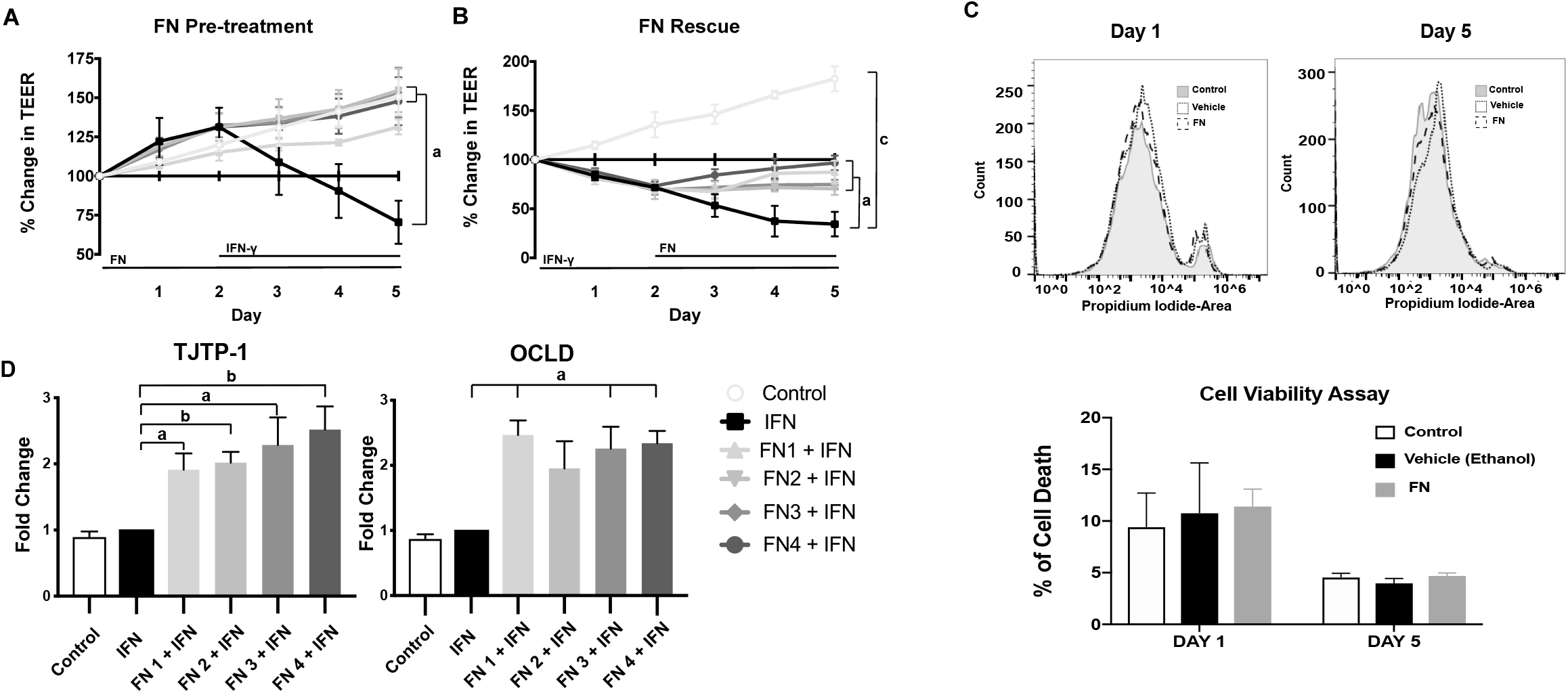
*F. vulgare* (FN) treatment of T84 cells improves tight junction functionality. A) FN pretreatment of T84 cells treated with IFN-γ prevents decrease of TEER. B) FN addition post IFN-γ treatment shows restorative effects on TEER. C) Propidium Iodide staining of Control, FN, and Vehicle treated T84 cells show no change in cell viability. D) FN treatments increase transcription of tight junction genes TJTP-1 and OCLD. N=4; a = P < 0.05; b = P < 0.01; c = P < 0.001; FN 1-FN 4 correlate to increasing doses of *F. vulgare*. Data shown as mean + SD. Significance calculated against IFN-γ group in all experiments.

To further elucidate the beneficial effects of *F. vulgare* on barrier function, expression of mRNA for selected tight junction proteins was evaluated via qRT-PCR. TJTP-1 and OCLD were examined as a possible explanation for the ability of *F. vulgare* extract to protect barrier function. There were significant increases in mRNA for both TJTP-1 and OCLD in *F. vulgare-treated* groups compared to the IFN-γ only treated groups (Figure 1D, P<0.03). TJTP-1 transcripts increased with rising *F. vulgare* dosages, with the highest concentration of *F. vulgare* eliciting a 2.2 fold increase compared IFN-γ treated cells. There was also a significant increase in TJTP-1 between FN 1 and FN 4 dosages. OCLD transcripts were increased by *F. vulgare* extract between 1.9 and 2.5 fold across all conditions. This effect was not dependent on the concentration of *F. vulgare* extract, unlike TJTP-1 expression.

### *F. vulgare* attenuates activation of STAT1

STAT1 activation was examined to determine other possible effects of *F. vulgare* on epithelial cells. Western blotting for activated phosphorylated STAT-1 (pSTAT-1) showed marked decreases in each of the *F. vulgare* treated groups, with the control group showing little pSTAT-1 as expected (Figure 2). Analysis of the ratio between pSTAT-1/STAT-1 showed a significant dose-dependent reduction in the pSTAT-1/STAT-1 ratio in response to *F. vulgare*. The lowest dosage of *F. vulgare* (4.5μL/mL) did not significantly reduce the pSTAT-1/STAT-1 ratio, while the three higher doses did reduce pSTAT-1/STAT-1 levels. The largest dose (9.0μL/mL) elicited a 76% reduction in pSTAT-1/STAT-1 based on the calculated band densitometry (Figure 2, P < 0.03).

**Figure 2.**
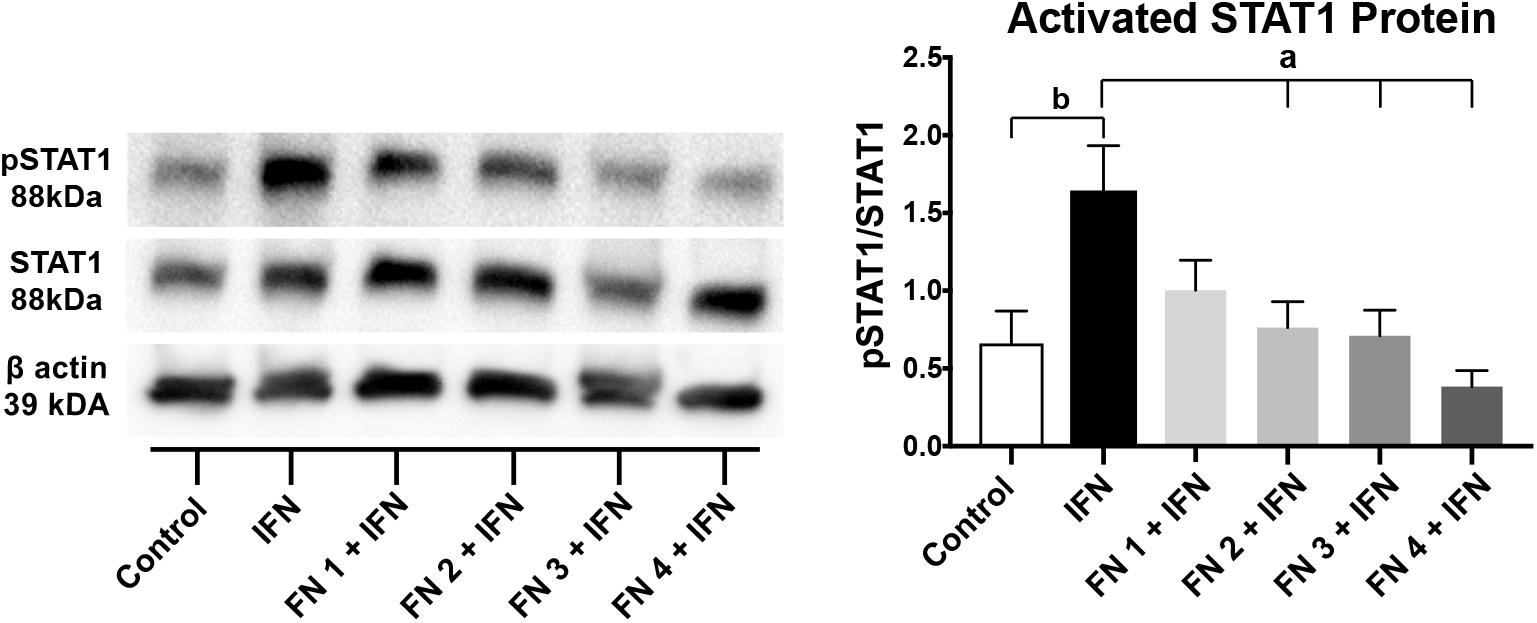
*F. vulgare* treated T84 cells show decreased STAT1 activation. Western blot analysis of pSTAT1 and STAT1 show decreasing pSTAT1 with increasing doses of *F. vulgare*. N=3; a = P < 0.05; b = P < 0.01; c = P < 0.001; FN 1-FN 4 correlate to increasing doses of *F.vulgare*. Data shown as mean + SD. Significance calculated against IFN-γ group in all experiments.

### *F. vulgare* exerts protective effects on barrier integrity in DSS colitis

To investigate whether our findings were relevant *in vivo*, parameters of colitis and barrier integrity were measured in mouse intestinal tissues following induction of colitis in the presence or absence of treatment with the *F. vulgare* extract. Upon sacrifice, 3 of 4 DSS treated mice presented with bloody stool, while the mice treated with *F. vulgare* had softer and lighter colored stool samples. Colonic tissue obtained from DSS treated mice was also more friable, or easier to damage through manipulation. *F. vulgare* treated and control mice demonstrated normal colonic integrity when handled. The TEER across mid colonic tissues of *F. vulgare* treated mice showed significantly higher values than DSS treated mice (Figure 3A, n=4). There was no observed difference among control mice and *F. vulgare* treated mice.

**Figure 3.**
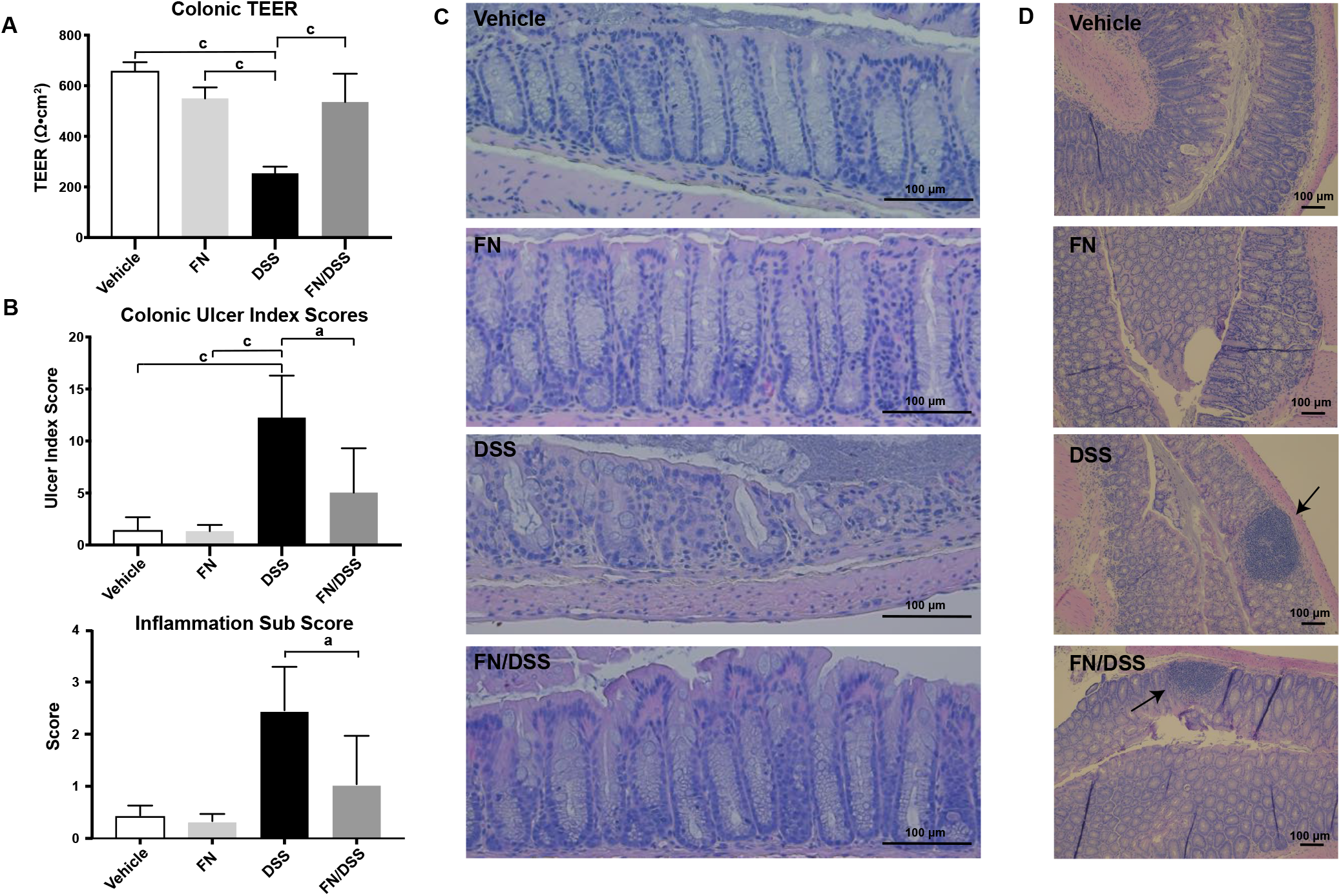
*F. vulgare* has beneficial effects on barrier integrity in mice with DSS colitis. A) *F. vulgare* prevents loss of TEER in FN/DSS treated mice compared to DSS treated mice. B) *F. vulgare* treated mice show improved ulcer indices compared to DSS treated mice. Inflammation sub score also shown, mirror improvement in overall ulcer indices. C) Representative H&E staining of mid colons of treated mice showing reduced inflammation, ulceration and architectural changes in *F. vulgare* treated mice. D) Representative H&E staining of mid colon demonstrating inflammation in DSS and *F. vulgare/DSS* treated mice. N=4; Figure 3C: 20x magnification, Figure 3D: 10x magnification; a = P < 0.05; c = P < 0.001

Histological analysis of colonic mucosa revealed decreases in crypt distortion, depth of ulceration and inflammatory infiltrates in DSS/*F. vulgare* mice vs. DSS mice while the histologic appearance of colons from control and *F. vulgare* groups were normal. Inflammation subscores confirmed the potency of DSS treatment, showing a significant inflammatory response in DSS treated mice and a lesser response in *F. vulgare/DSS* treated mice (Figure 3B, E). Ulceration was present in 50% of DSS mice, with a representative ulcer shown (Figure 3C). Ulcer indices were decreased in *F. vulgare/DSS* mice vs DSS mice (Figure 3B, 3C, P < 0.001).

### *F. vulgare* attenuates activation of STAT1 in DSS mice

The level of pSTAT1 and total STAT1 protein was measured in mouse colonic tissues from control, *F. vulgare*, DSS and FN*/*DSS groups. Western blot analysis showed that *F. vulgare/DSS* treated mice had significantly decreased levels of pSTAT1 compared to DSS treated mice (Figure 4).

**Figure 4.**
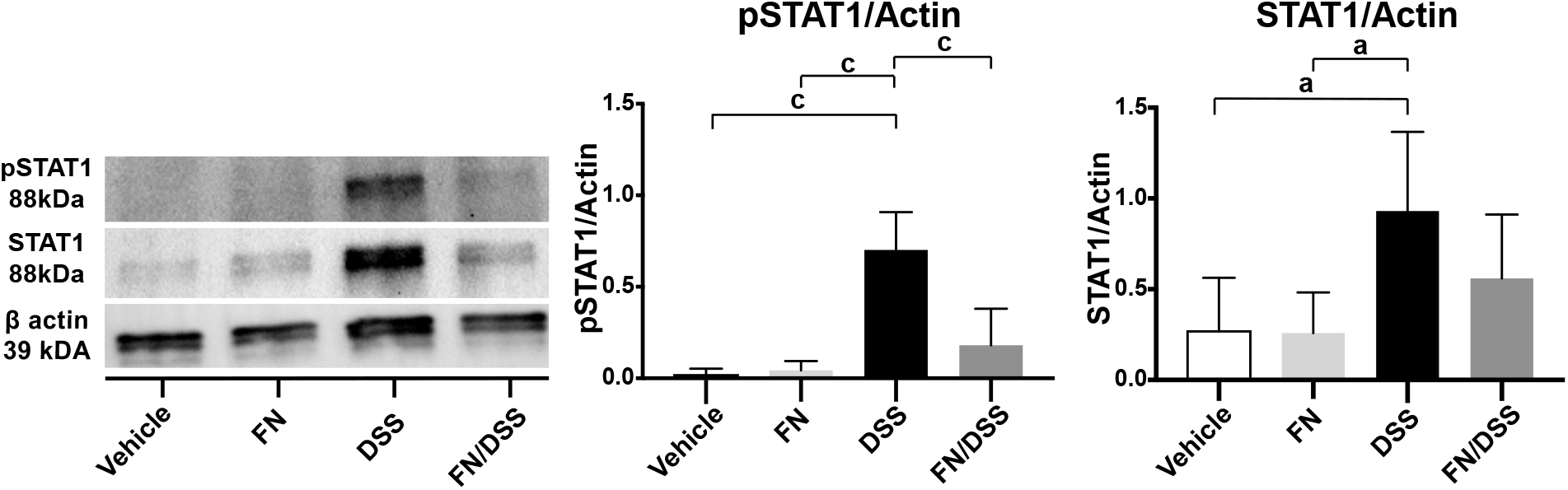
*F. vulgare* reduces protein expression of pSTAT1 in mice with DSS colitis. Western blot analysis demonstrates significantly reduced pSTAT1 in FN/DSS mice compared to DSS mice. Quantification of band density shown. pSTAT1 and STAT1 density adjusted for β-actin levels. Data presented as mean + upper SD. N=4; a = P < 0.05; c = P < 0.001

As expected, minimal pSTAT1 was detected in control and *F. vulgare* mice, indicating no active inflammatory processes (Figure 4). Treatment with DSS increased levels of total STAT1 compared to controls and *F. vulgare* did not significantly attenuate this increase (Figure 4). pSTAT1 and STAT1 levels were compared to actin levels due to changes in membrane background preventing direct comparison of pSTAT1 to STAT1 band density.

## Discussion

In this study, an oil extract of *F. vulgare* was shown to downregulate the STAT1 activation and improve gastrointestinal barrier function in both *in vitro* and *in vivo* models. *F. vulgare* seeds are known to contain volatile compounds and odorants including biologically active anethole.[41, 42] Previous studies have shown that *F. vulgare* has anti-oxidant effects and it is used in some areas of the world as a complementary and alternative treatment for IBD, but the effect of *F. vulgare* on barrier function and the JAK/STAT pathway, a major contributor to IBD pathogenesis, had not previously been examined.[9] The current medical treatment of IBD includes aminosalicylates, glucocorticosteroids, immunosuppressants, and biological therapies which all have side effects ranging from mild to life-threatening.[43, 44] Due to the common side effects of current IBD treatment alternative therapies, such as *F. vulgare*, have potential adjunctive roles.

This work investigated the effect of *F. vulgare* on tight junction proteins in the intestinal epithelium. The T84 cell line was used as a simple *in vitro* model reflective of colonic epithelial properties and treated with IFN-γ to model inflammation. T84 cells that were pre-treated with *F. vulgare* prior to IFN-γ treatment displayed significantly increased transcription of OCLD and TJPT-1 compared to controls that did not receive pre-treatment. Most notably, increases in transcription of TJTP-1 showed a dosage effect with increased transcription seen as the dose of *F. vulgare* was increased. OCLD and TJTP-1 are involved in regulating tissue electrical resistance, evidenced previously by OCLD-dependent increases in TEER.[45] Increased TEER and improved cell-cell interactions have been linked, which is largely due to strengthening of the zonula occludens junction, where OCLD and TJTP-1 are located. Epithelia with high TEER have been shown to have zonula occludens with more strands compared to epithelia with low TEER.[33] The increases in OCLD and TJTP-1 transcripts provide a potential explanation for the significantly improved TEER under inflammatory conditions. Interestingly, our data did not show the expected decreases in OCLD nor TJTP-1 mRNA in the IFN-γ only treatment group that have been established in previous literature.[46] Instead, similar levels of OCLD and TJTP-1 mRNA were noted between the IFN-γ treated and control cells. The absent tight junction modulation is not attributed to lack of response to IFN-γ challenge, as evidenced by the increase in pSTAT1 levels. Previous studies have shown tight junction regulation can be controlled via intracellular trafficking of tight junction associated proteins. TNF cytokines have been shown to induce calveolin-1 mediated endocytosis of occludin and restructuring of ZO-1 structures. [47, 48] Therefore, potentially tight junction modulation is occurring at the level of protein (intracellular trafficking) and not at the level of transcript. Our data suggest that *F. vulgare* attenuates the reduction of important barrier proteins in a model of the inflammatory state *in vitro*, with evidence of similar effects in the *in vivo* murine model. Studies also revealed several mechanisms for cytokines to disrupt the apical junction complex including affecting peri junctional cytoskeleton involving Myosin regulatory light chain [49, 50], disrupting apical actin [51], and disrupting lipid composition [52]. It is also reported that the proapoptotic effect of cytokines is independent of their influence on the epithelial junction complex [53].

In the DSS colitis mouse model, *F. vulgare* was shown to ameliorate structural pathology and protect against DSS-induced TEER loss. DSS-induced colitis has previously been shown to involve macroscopic damage such as multifocal gland dilation, epithelial erosion and infiltration of inflammatory cells into the lamina propria.[39, 54, 55] In this study, the mid colon of mice was analyzed for the severity of ulcers and tissue damage. A significantly decreased ulcer index was found in the mid colon from *F. vulgare/DSS* mice compared to DSS mice. Additionally, our finding that *F. vulgare* treatment prevented loss of TEER in DSS mice also presents evidence pointing toward an ameliorative role. Thus, we conclude that *F. vulgare* mitigates colitis symptoms at both the macroscopic and histological level.

Based on the protective effect of *F. vulgare* on barrier function and integrity during inflammation, the JAK/STAT pathway was investigated. Western blotting demonstrated levels of total STAT1 to be similar across experimental conditions. STAT1 is a major mediator in the JAK/STAT pathway leading to downstream transcription of inflammatory gene targets.[56] Increased protein expression of STAT1 has been seen in ulcerative colitis in humans.[57] Despite this, *F. vulgare* significantly attenuated the ability of IFN-γ to elevate pSTAT1 relative to total STAT1levels, suggesting that STAT1 activation and downstream events were impaired (Figure 5).

**Figure 5.**
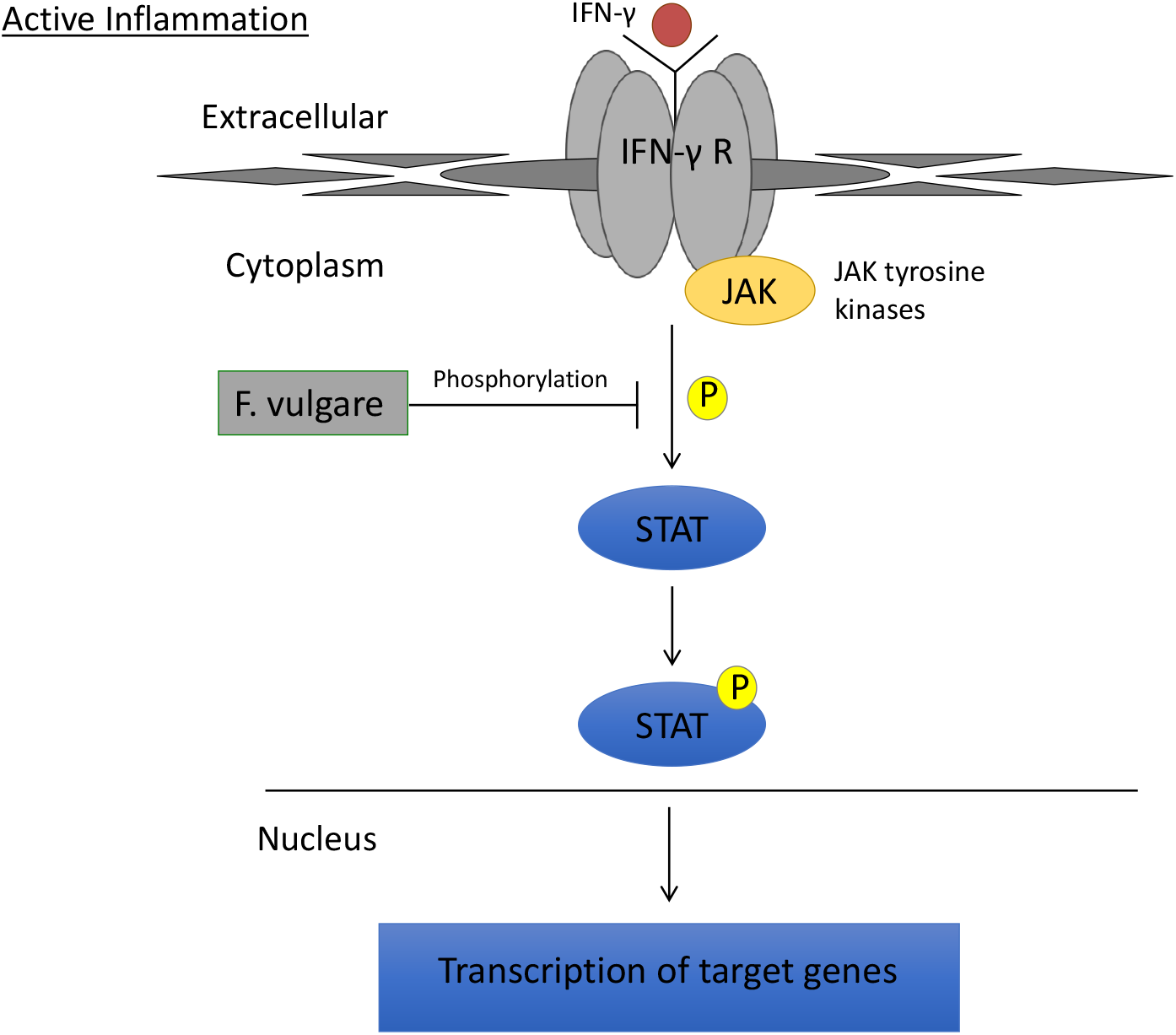
Schematic representation of mechanism of STAT pathway attenuation in T84 model. F. vulgare prevents phosphorylation of STAT1, preventing transcription of other inflammatory genes.

In murine mid colon, pSTAT1 was elevated when colitis was induced by DSS, as it is in the setting of human IBD. pSTAT1 was significantly decreased in *F. vulgare/DSS* mice vs. mice that received DSS alone. Therefore, *F. vulgare* is likely reducing activation of the STAT1 primarily by inhibiting phosphorylation of STAT1, as it did *in vitro* (Figure 4). We note that there was also a nonsignificant decrease in total STAT1 relative to actin in *F. vulgare*/DSS mice vs. DSS mice. Thus, comparing levels of pSTAT1/STAT1 as a ratio appeared to mask the absolute level of inflammatory response generated within the mice. Importantly, mice treated with only *F. vulgare* had similar expression of STAT1 to that in controls. This suggests *F. vulgare* plays a role in the state of active inflammation *in vivo*, inhibiting function of STAT1 only when mice are exposed to DSS. This may be similar to the known protection of epithelial barrier function by the Crohn’s disease associated gene protein tyrosine phosphatase.[58] Additionally, STAT phosphorylation has been shown to decrease TEER via Claudin-2 induction and prevent expression of adipogenesis related genes, both of which were also shown to be reversible via JAK inhibitor treatment, thereby reducing STAT phosphorylation.[59, 60]

Due to the success of *F. vulgare* in mitigating inflammatory response in both models of IBD, *F. vulgare* appears to be a promising candidate for further clinical trials to measure its efficacy as a complementary or alternative therapeutic. The improvements in gut health seen in this manuscript and other manuscripts examining herbal remedies lend credence to the notion of many undiscovered remedies or therapies for other chronic diseases. Identification of the active component within fennel seed extract was not explored in this study, limiting our understanding of the exact mechanisms of action. Future studies could pursue this knowledge, which would yield a more controllable therapeutic. The human equivalent dosage of *F. vulgare* extract given to mice in this study is 146 mg/kg; therefore interested clinical trials could consider starting dosages of 14.6 mg/kg. [61] Extract could be delivered in a similar way as given to mice (using water as vehicle) or simply added to food until target dosage is reached.

## Conclusion

Our findings regarding *F. vulgare* and its effect on the STAT pathway show parallels to current therapeutic approaches for IBD. Systemic glucocorticoids, often used to induce remission of IBD, lead to decreased levels of pSTAT1 in mucosal samples of patients with ulcerative colitis.[57] A novel JAK inhibitor, tofacitinib, was recently shown to induce clinical remission of UC in a randomized phase 3 trial.[62, 63] The treatment shown here with *F. vulgare* is in line with these IBD treatment modalities as all three target the JAK/STAT pathway to critically reduce inflammatory signaling in the intestines.

Due to the protective role of *F. vulgare* on barrier function and inflammatory proteins, *F. vulgare* has a potential role in IBD treatment. *F. vulgare* reduced histological and functional damage in mice with DSS-induced colitis. *F. vulgare* was also shown to have an anti-inflammatory effect through reduced activation of the STAT pathway in mice with DSS-induced colitis and in intestinal epithelial cells. As therapeutics targeting JAK/STAT activity are showing promise in the treatment of IBD, this further underscores the potential for *F. vulgare* in combatting disease manifestations. Additionally, such studies lend credence to the notion of consuming fennel seeds for digestive health.

## Abbreviations

IBD: inflammatory bowel disease
IBS: irritable bowel syndrome
HE: hematoxylin and eosin
JAK: Janus Kinase
STAT: Signal Transducer and Activator of Transcription
IFN-γ: interferon-gamma
pSTAT1: phosphorylated STAT1
*F. vulgare*: Foeniculum vulgare
TJP-1: tight junction protein-1
DSS: dextran sodium sulfate
TEER: transepithelial electrical resistance

## Declarations

### Ethics approval and consent to participate

Animal experiments were conducted per international guidelines. The protocols of this animal research study were approved by the UCSD IACUC. No animals died during the experiment, except by humane euthanasia.

### Consent for Publication

Not applicable.

### Funding

The authors extend appreciation to CCFA for summer award.

### Availability of data and materials

All data generated or analyzed during this study are included in this published article.

### Competing interests

The authors declare that they have no competing interests.

## Acknowledgments

The authors would like to extend their appreciation to Hal Hoffman and Lance Prince for helpful discussions.

## Authors’ Contributors

JR, PK, TL, ND executed experiments, performed data analysis and initial drafting of manuscript. MDM, KO, BD assisted in murine studies. JR, KO, BD, MS, JBL performed blinded tissue analysis. KEB, RM and MS contributed to interpretation of the data. MS coordinated the group, research and manuscript. All authors have been involved in the drafting, critical revision and final approval of the manuscript for publication.

## Notes

### Competing Interest Statement

The authors have declared no competing interest.

